# MetaCarto: biologically faithful automatic layout for genome-scale metabolic maps

**DOI:** 10.64898/2026.09.19.752882

**Authors:** Tianyu Wu

## Abstract

**Motivation:** Genome-scale metabolic reconstructions containing thousands of reactions are now curated and published for many organisms, but their visualization still relies heavily on manually constructed pathway maps. As these models grow in size and number, an automatic method is needed to organize their reactions into readable maps for visualization and downstream analysis.

**Results:** We developed MetaCarto, an automatic layout framework for genome-scale metabolic maps. MetaCarto identifies a biologically meaningful primary connection for each reaction and then organizes these connections into readable pathway-scale layouts, separating the biological representation of a reaction from its geometric placement. The resulting maps show strong agreement with curated KEGG pathway representations while maintaining geometric quality comparable to established graph-layout approaches. We applied MetaCarto to all 108 BiGG models and generated 2621 pathway maps without manual rearrangement. As a large-scale example, the human Recon3D reconstruction, containing 10 600 reactions, was automatically decomposed into 93 maps and laid out in 107 s. The generated maps are exported as Escher-compatible JSON and SVG files and can be directly browsed, edited, and used for metabolic data visualization.

**Availability and implementation:** Source code is available under CC BY 4.0 at github.com/forxhunter/MetaCarto. The complete collection of generated maps is available at GitHub map collection and can be browsed interactively at forxhunter.github.io/escher.

**Supplementary information:** Supplementary data are available at github.com/forxhunter/MetaCarto.

## 1 Introduction

Genome-scale metabolic reconstructions describe the metabolism of an organism as networks containing thousands of reactions. The *Escherichia coli* model iJO1366 contains 2583 reactions, and the human reconstruction Recon3D contains 10 600 (Orth *et al*., 2011; Brunk *et al*., 2018); the BiGG database currently distributes 108 such models (Norsigian *et al*., 2019). Researchers interpret these networks through pathway maps, which place reactions, fluxes, and molecular measurements into a biochemical context that a stoichiometric matrix alone does not provide. Yet the pathway maps most commonly used for this purpose, including KEGG diagrams and Escher maps, are largely constructed by experts (Kanehisa, 2000; King *et al*., 2015). Manual drawing does not scale easily to models with thousands of reactions, many organisms, and reconstructions that are continually revised. Automatic generation of readable metabolic maps is therefore increasingly important. Most work on automatic metabolic-network visualization has treated this as a graph-layout problem. Methods have been developed for organism-scale overviews, pathway-preserving layouts, genome-scale drawing, interactive visualization, and scalable pathway organization (Karp *et al*., 2022; Bourqui *et al*., 2007; Lambert *et al*., 2011; Jensen and Papin, 2014; Kelley *et al*., 2017; Moškon *et al*., 2018; Chazalviel *et al*., 2018; Wu *et al*., 2019; Aichem *et al*., 2021; Krishnamurthy *et al*., 2026). Their quality is consequently evaluated mainly through geometric properties such as edge crossings, bends, node separation, density, and orthogonality (Purchase, 1997; Bourqui *et al*., 2007; Schreiber *et al*., 2009; Lambert *et al*., 2011; Wu *et al*., 2019). This framing carries an implicit assumption: if an automatic layout produces a well-organized graph, it has produced a good metabolic map.

Metabolic reactions expose a problem with this assumption. A reaction usually contains several substrates and products, but a pathway map must decide which relationship forms the main pathway connection. Pyruvate dehydrogenase, for example, converts pyruvate and CoA to acetyl-CoA, CO_2_, and NADH. It can be drawn along the carbon transformation, pyruvate →acetyl-CoA; along the carrier transformation, CoA →acetyl-CoA; or along the redox pair, NAD^+^ →NADH. Each representation can occupy exactly the same amount of space and have the same number of crossings, bends, and aligned edges. Yet they place the reaction on different biochemical backbones. Geometry can determine where an edge is drawn, but it cannot determine which edge should represent the reaction.

This leads to the central question of this work: *does geometric quality indicate that an automatically generated metabolic map is biologically correct?* To answer it, we separate these two properties experimentally. We use curated KEGG pathway files as a reference for the substrate–product connections drawn by human curators, providing a direct measure of biological agreement. We then perturb the biological rule used to choose the primary connection of each reaction while leaving the downstream layout procedure unchanged. The resulting maps remain geometrically similar even when their agreement with curated pathway connections decreases substantially. Thus, geometric quality and biological correctness are distinct properties of a metabolic map. A layout can look well organized while placing reactions on the wrong biochemical backbone.

This distinction changes the order in which automatic metabolic maps should be constructed. We developed MetaCarto around this principle. For each reaction, MetaCarto first selects one primary substrate–product connection using conserved chemical composition together with a constraint that prevents ubiquitous carriers from dominating the pathway backbone. Other substrates and products are retained as side branches. Only after this biological representation has been chosen does the method determine reaction orientation and geometric placement. The reduced network is then arranged into pathway-scale maps using flux-guided orientation, explicit cycle placement, and layered graph layout (Lewis *et al*., 2010; Sugiyama *et al*., 1981; Brandes and Köpf, 2002). The resulting maps are exported directly as Escher-schema JSON and can therefore be opened, edited, and combined with experimental data in existing visualization tools.

Figure 1 illustrates the result for four pathways from the human Recon3D reconstruction. The examples show how reactions are organized around a primary biochemical backbone while cofactors and other secondary participants remain visible as side branches. Cyclic pathways are represented explicitly as cycles, and parallel pathway branches remain visually distinguishable without manual rearrangement.

**Fig 1.**
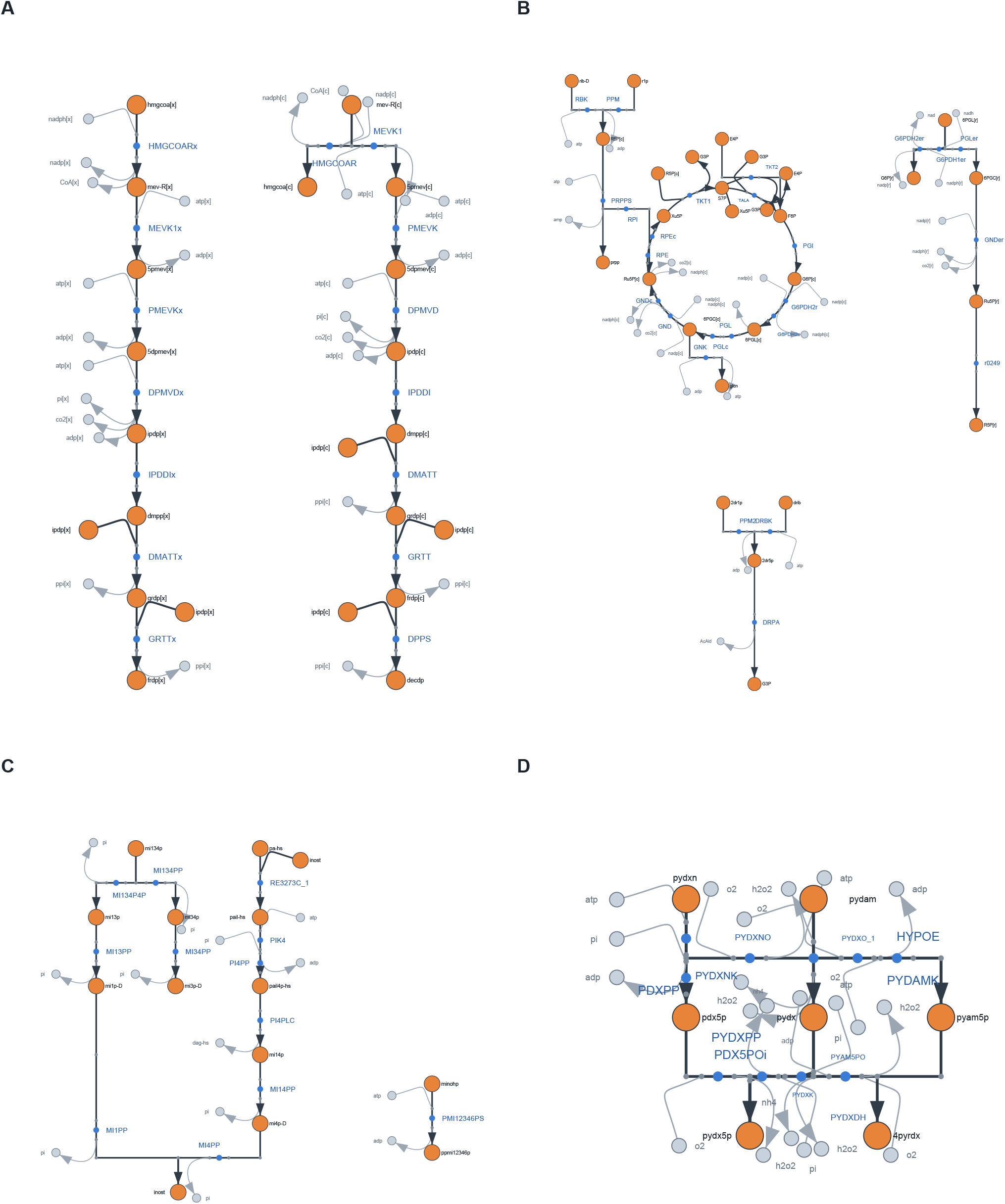
Four automatically generated pathway maps from the human Recon3D reconstruction, which contains 10 600 reactions. Each reaction is represented by one primary substrate–product connection forming the pathway backbone, while its remaining substrates and products are retained as side branches. Each panel corresponds to one pathway cluster (Section 2.1). (**A**) Terpenoid backbone biosynthesis, showing parallel peroxisomal and cytosolic branches. (**B**) Pentose phosphate pathway, with its cycle placed explicitly as a ring. (**C**) Inositol phosphate metabolism. (**D**) Vitamin B6 metabolism. All layouts were generated by MetaCarto without manual rearrangement.

We applied this framework to the complete BiGG collection. MetaCarto generated 2621 pathway maps covering 240 398 reactions across all 108 models without manual layout. The largest example, Recon3D, contains 10 600 reactions and was automatically decomposed into 93 pathway maps and processed in 107 s. The resulting collection provides a consistent visualization resource across the full model database. The broader conclusion is that automatic metabolic-map generation has two separate problems: deciding what biochemical relationship each reaction should represent and deciding where that representation should be placed. Graph-layout metrics evaluate the second problem but cannot validate the first. Treating biological representation as an explicit, independently measured step makes it possible to evaluate automatic metabolic maps on the property that ultimately determines what pathway a reader sees.

## 2 Inputs and evaluation

This section defines the model inputs, pathway-scale decomposition, and evaluation protocol used to develop and test MetaCarto. The algorithm itself is described in the following section.

### 2.1 Input models and map decomposition

MetaCarto reads genome-scale metabolic models in BiGG SBML or JSON format using COBRApy (Ebrahim *et al*., 2013). We applied it to all 108 BiGG models, ranging from the 95-reaction *E. coli* core model to the 10 600-reaction human Recon3D model. The required inputs are reaction stoichiometry, metabolite molecular formulae, and reaction reversibility; flux bounds are additionally used for reaction orientation (Section 3.2).

Genome-scale models are divided into pathway-scale maps before layout. Reactions are grouped using model`subsystem` annotations where available, followed by KEGG pathway annotations. Reactions assigned by neither source are grouped by greedy modularity after removing currency metabolites (Clauset *et al*., 2004; Hagberg *et al*., 2008). Initial clusters contain 6–60 reactions; clusters belonging to the same KEGG BRITE metabolic function are then merged up to 120 reactions. This decomposition determines which reactions are drawn together but does not otherwise change the layout algorithm.

Each reaction is represented in the layout by one primary substrate–product connection, with its remaining participants retained as side branches. The rule for selecting this connection is described in Section 3.1. Generated maps are exported as Escher-compatible JSON (King *et al*., 2015) and can also be rendered as SVG or vector PDF.

### 2.2 Evaluation protocol and metrics

We evaluate MetaCarto using two complementary measurements: agreement between the selected primary connection and curated pathway representations, and the geometric quality of the resulting map.

#### Curated-connection agreement

KEGG KGML files record the substrate and product entries connected through each displayed reaction (Kanehisa, 2000). Our reference collection contains 1010 organism-specific pathway files from human, mouse, *S. cerevisiae*, and *E. coli*. KGML does not explicitly designate one substrate–product pair as the unique main connection. For a KEGG reaction *R*, we therefore define the reference set as all connections drawn for that reaction,

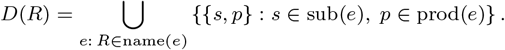

The pair {*s, p*} is treated as unordered because connection identity and reaction orientation are evaluated separately. A BiGG reaction is counted as agreeing with the reference when its selected substrate–product pair, mapped through`kegg.reaction` and `kegg.compound` annotations, belongs to *D*(*R*). Reactions lacking the required mappings are excluded from this evaluation but are still drawn.

#### Geometric quality

Each generated map is evaluated using standard graph-layout and readability metrics: axis-aligned edge fraction, crossings per edge, longest straight run, minimum node separation, label–label, label–node, and label–edge overlap, local density, bounding-box occupancy, and aspect ratio. We additionally estimate print legibility by scaling each map to a 180 mm by 240 mm page and measuring the resulting metabolite-label size.

For comparison with curated layouts, KEGG pathway coordinates are converted to an Escher-like representation while preserving their original positions. After deduplication by pathway number, 117 distinct KEGG layouts are available for geometric comparison. Because label-overlap measures depend on notation and minimum node separation uses Escher-specific normalization, direct KEGG–MetaCarto comparisons use crossings per edge, local density, and aspect ratio.

#### Quality-control thresholds

For corpus-level quality control, each geometric metric is assigned a pre-specified acceptance band (Table 1). These engineering thresholds were fixed before generation of the full map collection and were not fitted to the resulting maps. The corresponding distributions from curated KEGG layouts are reported alongside them where the metrics are directly comparable.

**Table 1.**
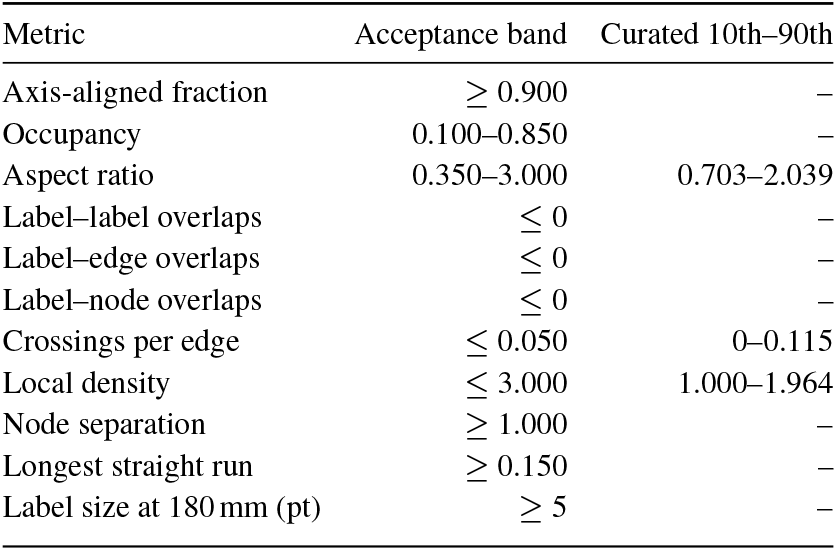
Pre-specified geometric quality criteria and the 10th–90th percentile range of the 117 curated KEGG layouts for directly comparable metrics. Acceptance bands were fixed before generation of the full map collection and were not calibrated to the generated maps.

| Metric | Acceptance band | Curated 10th–90th |
| --- | --- | --- |
| Axis-aligned fraction | $\geq 0.900$ | – |
| Occupancy | 0.100–0.850 | – |
| Aspect ratio | 0.350–3.000 | 0.703–2.039 |
| Label–label overlaps | $\leq 0$ | – |
| Label–edge overlaps | $\leq 0$ | – |
| Label–node overlaps | $\leq 0$ | – |
| Crossings per edge | $\leq 0.050$ | 0–0.115 |
| Local density | $\leq 3.000$ | 1.000–1.964 |
| Node separation | $\geq 1.000$ | – |
| Longest straight run | $\geq 0.150$ | – |
| Label size at 180 mm (pt) | $\geq 5$ | – |

## 3 Map construction and matched perturbations

MetaCarto constructs a metabolic map in two conceptually separate phases. The first determines *what each reaction represents* by selecting one substrate–product connection as its primary biochemical connection. The second determines *where that representation is placed* by orienting, arranging, packing, and rendering the resulting network.

This separation mirrors the central question of the study. Biological representation is evaluated by curated-connection agreement (Section 2.2),whereas geometric placement is evaluated by the layout metrics defined above. To test whether these two properties track one another, we perturb only the rule used to select the primary connection, leave all downstream layout steps unchanged, and redraw the same pathway clusters.

Matched geometric comparisons are performed cluster by cluster. *P* - values are Holm-corrected within each model over the complete set of comparisons (Holm, 1979), and matched-pairs rank-biserial correlations are reported as effect sizes (Wilcoxon, 1945; Kerby, 2014).

### 3.1 Biological representation and matched perturbations

A biochemical reaction can contain several substrates and several products. In a conventional bipartite reaction graph, a reaction is connected to all of these participants simultaneously. A pathway map, however, usually emphasizes one substrate–product relationship as the continuation of the pathway. MetaCarto represents each reaction by one *primary connection*, a single substrate–product pair that forms the pathway backbone. All other substrates and products are retained as side participants and remain visible in the final map. Selecting this pair determines which biochemical transformation the reaction contributes to the pathway. Candidate substrate–product pairs are first scored by heavy-atom composition overlap: the number of non-hydrogen atoms that can be matched by element, normalized by the heavy-atom count of the smaller metabolite. This provides a simple estimate of molecular continuity from molecular formulae alone and does not require atom mapping.

Composition overlap alone can favor large conserved carrier moieties. In pyruvate dehydrogenase, for example, CoA–acetyl-CoA can score above the pyruvate–acetyl-CoA connection even though the latter better represents the carbon pathway. Network connectivity does not reliably resolve this ambiguity because many central pathway metabolites are themselves highly connected. MetaCarto applies a carrier constraint before ranking candidate pairs. If at least one candidate avoids a metabolite in the curated carrier list, pairs involving that carrier are excluded from primary-connection selection. The remaining pairs are ranked by composition overlap with a connectivity damping term. When every candidate contains a carrier, no pair is excluded and composition overlap determines the selection. The resulting primary connection is evaluated against the curated reference *D*(*R*) defined in Section 2.2. This comparison is made before any coordinates are assigned, so curated-connection agreement measures the biochemical representation independently of geometric placement.

To test whether geometric quality reflects this biological choice, we define matched perturbations at the same selection stage. The first removes the carrier constraint while retaining the same composition-overlap scoring procedure, allowing carrier-containing pairs to compete directly with all other candidates. A stronger perturbation removes explicit cofactor handling altogether. In both cases, all downstream orientation, placement, packing, and rendering procedures remain unchanged. The same pathway clusters are redrawn after changing only the rule that determines which connection represents each reaction. Curated-connection agreement and geometric metrics can then be measured on matched maps. This design directly tests whether changing the biochemical backbone produces a corresponding change in conventional measures of layout quality.

### 3.2 Geometric layout and rendering

Once a primary connection has been selected for every reaction, all remaining steps operate on the reduced compound graph. They can change the direction, position, and appearance of a connection, but they cannot change which substrate–product pair represents the reaction.

#### Orientation

Reaction direction in a reconstruction does not always correspond to the direction in which neighbouring reactions should be read as a pathway. Reversible reactions may be stored in opposite orientations even when their net flux proceeds along the same biochemical route. MetaCarto therefore orients each primary connection using the sign of its parsimonious flux-balance-analysis flux (Lewis *et al*., 2010). When the calculated flux is zero, orientation falls back to the surrounding graph topology. Antiparallel reaction pairs are aligned along the same pathway axis instead of being represented as artificial two-node cycles.

#### Cycle placement

Directed cycles that remain after orientation are detected in the reduced compound graph and temporarily contracted into super-nodes before layout. After placement of the surrounding network, each cycle is expanded onto a ring and rotated so that its principal incoming connection faces the pathway that feeds it. This preserves recognizable cyclic pathway structure within an otherwise layered layout.

#### Layered placement

After orientation and cycle contraction, the remaining graph is converted to an acyclic representation using a greedy feedback-arc-set procedure (Eades *et al*., 1993). Nodes are assigned to layers and ordered within each layer to reduce crossings while preserving pathway structure (Sugiyama *et al*., 1981). Horizontal coordinates are assigned using the Brandes–Köpf method (Brandes and Köpf, 2002), producing long aligned pathway backbones. A separation pass is then applied to prevent overlaps and is repeated after side participants are inserted.

#### Component packing

A pathway map can contain several disconnected components. MetaCarto processes components from largest to smallest, using the larger components to establish the main layout and placing smaller components into remaining free regions on a grid. Packing changes the use of canvas space without altering the internal representation of any reaction.

#### Rendering

The final coordinates are converted to the Escher representation. Each primary connection is encoded as metabolite →multimarker →midmarker →multimarker →metabolite, with reaction direction determined by the corresponding stoichiometric coefficients. Substrates and products not selected as the primary connection are attached as short side branches. Paired cofactors such as ATP/ADP and NAD^+^/NADH are placed on the same side of the pathway where possible, and curved Bézier segments connect these secondary participants while keeping the primary pathway backbone visually continuous.

The final output is an Escher-schema JSON map in which the biochemical connection chosen for each reaction and the coordinates used to display it remain explicitly separable.

## 4 Implementation and benchmarks

MetaCarto is implemented in Python using COBRApy, NetworkX, and NumPy (Ebrahim *et al*., 2013; Hagberg *et al*., 2008). The implementation is deterministic and requires no stochastic optimization, random seed, or manual rearrangement; repeated runs on the same model produce identical Escher JSON.

### 4.1 Benchmark performance

#### Geometric placement

We compared MetaCarto with Graphviz DOT, NEATO, and FDP (Gansner and North, 2000), together with NetworkX spring and Kamada–Kawai layouts. All methods received the same reduced and oriented graphs and were evaluated with the same renderer and metric code. The benchmark contains 676 maps from eight organisms, with paired Wilcoxon signed-rank tests, Holm correction over 30 comparisons, and matched-pairs rank-biserial effect sizes (Table 2).

**Table 2.** Layout quality over 676 maps from eight organisms. Cells report matched-pairs rank-biserial effects relative to MetaCARTO; negative values favour MetaCsc>ARTO. P -values are Holm-corrected, and n.s. denotes P ≥ 0.05. The MetaCARTO column reports the median value for each metric.

| Metric | METACARTO | DOT | NEATO | FDP | spring | KK |
| --- | --- | --- | --- | --- | --- | --- |
| Crossings/edge | 0.000 | +0.93 | +0.83 | −0.18 | +0.66 | −0.69 |
| Axis-aligned | 0.615 | +0.67 | −1.00 | −0.99 | −0.99 | −0.99 |
| Local density | 1.838 | +0.22 | +0.19 | +0.35 | −0.74 | −0.37 |
| Aspect ratio | 0.838 | −0.67 | +0.57 | +0.77 | +0.57 | +0.34 |
| Occupancy | 0.420 | −0.85 | *n.s.* | +0.65 | *n.s.* | *n.s.* |
| Separation | 0.846 | *n.s.* | −0.41 | −0.46 | −0.31 | −0.86 |

Across six geometric metrics and five baselines, MetaCARTO performs comparably to DOT: each method is better on different aspects of the layout. The main separation is from force-directed methods. The median axis-aligned edge fraction is 0.615 for MetaCARTO, compared with 0.000–0.019 for NEATO, FDP, spring, and Kamada–Kawai, with rank-biserial effects of −0.99 or stronger and all *P <* 10−^15^.

#### Connectivity control

We also tested whether a purely connectivity-based rule could recover the same curated pathway connections without the carrier constraint. Following the compound-cloning strategy used by MetDraw (Jensen and Papin, 2014), metabolites above the 80th, 90th, or 95th connectivity percentile were duplicated before scoring against the curated reference.

Connectivity cloning recovers a median 95.7–96.1% of curated connections, but does so by retaining 1.75–2.86 candidate connections per reaction, giving 38.0–60.6% precision. MetaCARTO selects exactly one connection per reaction and reaches a median curated agreement of 94.8%. Thus connectivity cloning recovers the curated connection by retaining several alternatives, whereas MetaCARTO resolves the ambiguity to a single pathway connection.

### 4.2 Scale and released corpus

Runtime was measured on models spanning two orders of magnitude in reaction count (Table 4). The 10 600-reaction human Recon3D model is decomposed into 93 pathway maps and processed in 107 s: 64.2 s for decomposition and 32.2 s for compound-graph construction, orientation, placement, and rendering.

**Table 3.** Agreement with curated KEGG substrate–product pairs for MetaCARTO and connectivity cloning across three connectivity thresholds.

| Method | Recall | Precision | Edges/rxn |
| --- | --- | --- | --- |
| METACARTO | 94.8 | 94.8 | 1.00 |
| clone p80 | 96.1 | 38.0 | 2.86 |
| clone p90 | 95.7 | 48.2 | 2.19 |
| clone p95 | 96.1 | 60.6 | 1.75 |

**Table 4.** Wall-clock runtime in seconds for each processing stage.

| Model | Reactions | Maps | Load | Decompose | Draw |
| --- | --- | --- | --- | --- | --- |
| *E. coli* core | 95 | 3 | 0.0 | 0.1 | 0.1 |
| iYO844 | 1250 | 14 | 1.3 | 1.0 | 2.5 |
| iJO1366 | 2583 | 28 | 2.5 | 4.1 | 4.9 |
| Recon3D | 10 600 | 93 | 10.8 | 64.2 | 32.2 |

We applied MetaCarto to all 108 BiGG models, producing 2621 maps covering 240 398 reactions with no layout failure. The collection is released as Escher JSON and SVG and can be browsed through an Escher-based interface (Fig. 2).

**Fig 2.**
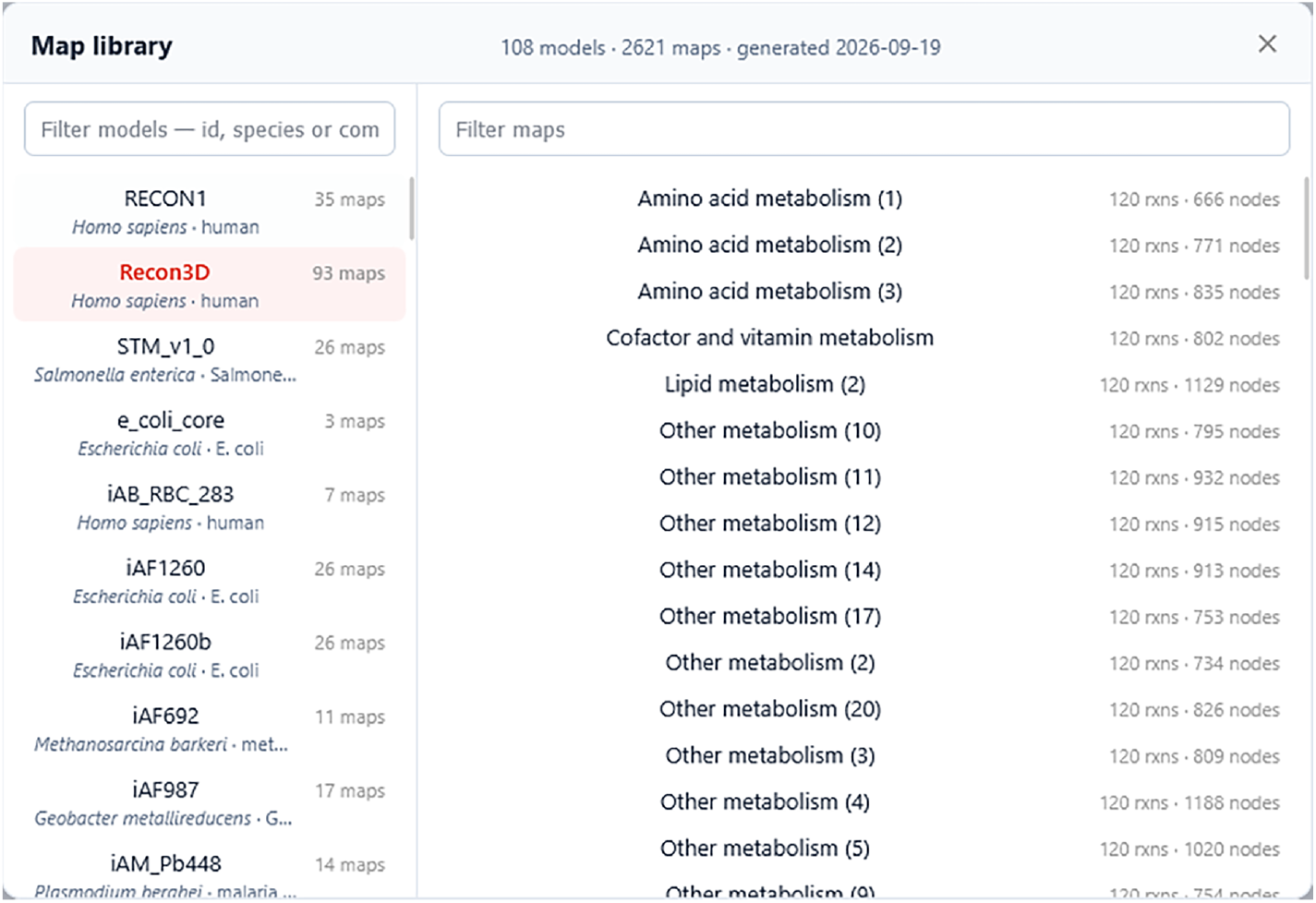
Browser interface for the released map collection. The library contains 2621 maps from all 108 BiGG models and can be searched by model identifier, species, strain, or common organism name. Individual pathway maps are listed with their reaction and node counts.

Across the full collection, the median axis-aligned edge fraction is 0.988, and 99.7% of maps satisfy the corresponding geometric acceptance criterion (Table 5). Occupancy and aspect ratio also remain within their pre-specified ranges for most maps. Crossings and local density are more variable, with medians of 0.056 per edge and 3.13, respectively. The main limitation appears at whole-model display scale: when maps are compressed to a 180 mm-wide page, the median metabolite label is only 1.9 pt. The released collection is therefore primarily suited to interactive navigation, whereas individual pathway-scale maps remain suitable for print.

**Table 5.** Quality-control metrics across all 2621 released maps. “In band” gives the fraction of maps satisfying the pre-specified criterion for each metric (Section 2.2 and Table 1).

| Metric | 10th | Median | 90th | In band (%) |
| --- | --- | --- | --- | --- |
| Axis-aligned fraction | 0.958 | 0.988 | 1.000 | 99.7 |
| Occupancy | 0.408 | 0.631 | 0.674 | 99.5 |
| Aspect ratio | 0.381 | 0.989 | 2.203 | 91.5 |
| Label–label overlaps | 0.000 | 0.000 | 1.000 | 86.5 |
| Label–edge overlaps | 0.000 | 0.000 | 3.000 | 76.7 |
| Label–node overlaps | 0.000 | 0.000 | 4.000 | 74.4 |
| Crossings per edge | 0.000 | 0.056 | 0.433 | 46.4 |
| Local density | 1.000 | 3.127 | 4.692 | 44.6 |
| Node separation | 0.367 | 0.367 | 1.444 | 18.8 |
| Longest straight run | 0.019 | 0.061 | 0.196 | 14.2 |
| Label size at 180 mm (pt) | 0.983 | 1.853 | 6.324 | 13.5 |

## 5 Discussion

Geometric quality and biological correctness are distinct properties of an automatic metabolic map. Removing MetaCarto’s carrier constraint changed which substrate–product connection represented a reaction while leaving the tested geometric metrics essentially unchanged, showing that crossings, spacing, orthogonality, and related layout measures cannot by themselves determine whether a reaction has been placed on the correct biochemical backbone. This explains why MetaCarto can have geometry comparable to Graphviz DOT while differing substantially in biological representation: geometry evaluates where a selected connection is placed, whereas curated-connection agreement evaluates whether the selected connection is appropriate. MetaCarto resolves this ambiguity by selecting one primary connection per reaction and reaches a median curated agreement of 94.8%, while connectivity-based cloning attains similar recall only by retaining multiple alternatives for the reader to resolve. Automatic metabolic-map methods should therefore report both biological representation and geometric quality. In practical terms, this separation makes it possible to generate consistent pathway maps at genome scale: MetaCarto produced 2621 maps covering all 108 BiGG models without manual layout, including the 10 600-reaction Recon3D reconstruction in 107 s.

### 5.1 Limitations

A practical limitation is that very large reconstructions cannot be displayed as a single fully readable Escher canvas because labels and reaction details become too small at whole-network scale. MetaCarto therefore decomposes large models into multiple pathway-scale maps, prioritizing readability and interactive navigation over a single giant canvas. In addition, although the generated layouts are evaluated systematically using curated KEGG connections and geometric metrics, most of the 2621 maps have not been individually reviewed by domain experts. Some local pathway assignments or visual arrangements may therefore still benefit from expert inspection and manual refinement for specific applications.

## Data availability

The MetaCarto source code is available under CC BY 4.0 at github.com/forxhunter/MetaCarto. The 2621 generated maps, each as Escher-schema JSON and as SVG, are available under the same licence at github.com/forxhunter/Awesome_visualization_Metabolic_Network and can be browsed at forxhunter.github.io/escher. JSON is what the viewer loads and SVG opens in any browser;`scripts/render_corpus.py` re-renders the collection to PNG or vector PDF from the same files.

The metabolic models are redistributed by BiGG Models (Norsigian *et al*., 2019) at bigg.ucsd.edu, and the curated reference layouts are KEGG KGML pathway files (Kanehisa, 2000) from kegg.jp/kegg/xml. No new experimental data were generated.

## Acknowledgements

An LLM-based coding assistant was used to help write and debug the software and to draft its code documentation, and to check the spelling and grammar of this manuscript. The author is responsible for the correctness of the code and of the text. Supplementary Section 1 gives the full declaration.

