## Supplementary File for "MetaCarto: biologically faithful automatic layout for genome-scale metabolic maps"

Tianyu Wu

#### 1 Declaration of the use of large language models

An LLM-based coding assistant (Claude, Anthropic) was used during the development of the MetaCarto software. Its use falls under the uses the Journal lists as acceptable, and was limited to the following.

- **Assistance with code writing and debugging.** The assistant proposed and reviewed implementation code and helped diagnose defects. One such defect is reported in §4 because it changed a result: an ablation in the benchmark harness was found to patch a constant that the layout code no longer read, so the variant it was supposed to measure was disabled in name only. The author is responsible for the correctness of all code.
- **Code documentation.** Module and function docstrings were drafted with assistance and checked by the author.
- **Spelling and grammar.** The manuscript text was checked for spelling and grammatical errors.

The algorithmic design, the choice of evaluation, the experiments, the interpretation of the results and the conclusions are the author's. No text, figure, table or reference in the manuscript was drafted by an LLM from a prompt.

#### 2 Data and software availability

- **Source code.** `https://github.com/forxhunter/MetaCarto`, CC BY 4.0.
- **Generated maps.** 2621 maps covering all 108 BiGG models, each published as Escher-schema JSON and as SVG: `https://github.com/forxhunter/Awesome_visualization_Metabolic_Network`, CC BY 4.0. JSON is what the Escher viewer loads and SVG opens in any browser; `scripts/render_corpus.py` re-renders the collection to PNG or vector PDF from the same files.
- **Viewer.** `https://forxhunter.github.io/escher/`.
- **Input models.** BiGG Models, `http://bigg.ucsd.edu/`.
- **Curated reference layouts.** KEGG KGML pathway files, `https://www.kegg.jp/kegg/xml/`.
- **Benchmark results.** Every number in the manuscript is read from a JSON file under `benchmarks/results/` in the source repository. Each file records the git commit that produced it.

### 3 Benchmark protocol

#### 3.1 Common adapter

Every method is scored through the same pipeline so that only coordinate assignment varies between them. For each cluster, MetaCarto builds the compound graph and orients it; that graph is handed to the placement method; the returned coordinates are normalised to a common pitch; and the result is rendered to Escher JSON by the same renderer and scored by the same metric code.

Two consequences follow from holding the pipeline fixed.

- The baselines receive MetaCarto's primary-compound reduction. MetDraw does not perform this reduction, so the Graphviz baselines here are drawing a sparser and easier graph than MetDraw would.
- MetaCarto is measured with straight edges rather than its own orthogonal router, because the router is applied after placement and would be credited to whichever method it ran on. Routing a Kamada–Kawai drawing through it produced a layout scoring 0.969 axis-aligned, measuring the router rather than the placement.

#### 3.2 Statistics

Each metric is compared per map between MetaCarto and one baseline by a paired Wilcoxon signed-rank test. Effect sizes are matched-pairs rank-biserial correlations; Cliff's delta is not used because the samples are paired. $P$-values are Holm-corrected over the whole family of 30 tests ($6$ metrics $\times$ $5$ baselines), not per metric. Pairs in which both methods give an identical value carry no rank and are excluded, which is why the number of pairs used differs between metrics.

#### 3.3 Corpus

676 maps drawn from eight models: RECON1, iAF692, iCHOv1, iJO1366, iMM904, iNJ661, iSB619 and iYO844.

### 4 The ablation harness defect

The ablation that removes the curated-carrier tier originally patched a constant, `_COFACTOR_CUTOFF`, which had been the tier threshold when cofactor-ness was a score cutoff. The tier was later changed to membership of a curated carrier list, and nothing read the constant afterwards. The patch was therefore inert, and the ablation reported agreement with KEGG identical to the full method — to the individual reaction — on two models, which reads as the carrier constraint being worth nothing.

Corrected so that the ablation empties the curated list, the constraint is worth a median 5.1 percentage points of agreement across the corpus. The companion ablation, which was intended to remove tiering and damping together, had been removing only the damping; corrected, it costs 35.5 points pooled. The dead constant has been deleted rather than left in place, and a regression test asserts that the ablation changes the pair chosen for pyruvate dehydrogenase before its number is reported.

### 5 Full ablation results

Values are the median of the ablated variant over the clusters of that model; * marks $P < 0.05$ after Holm correction over the 30-test family.

| Variant / metric | iJO1366 | iMM904 | iNJ661 | iYO844 |
|---|---|---|---|---|
| no_bk | | | | |
| Crossings/edge | 0.000 | 0.000 | 0.000 | 0.000 |
| Axis-aligned | 0.263* | 0.286* | 0.432* | 0.300* |
| Local density | 1.860 | 1.000 | 1.810 | 1.000 |
| Aspect ratio | 0.838 | 0.802 | 0.773 | 0.754 |
| Separation | 0.846 | 0.733 | 0.757 | 0.812 |
| no_cofactor_handling | | | | |
| Crossings/edge | 0.000 | 0.000 | 0.000 | 0.000 |
| Axis-aligned | 0.719* | 0.710 | 0.742 | 0.714 |
| Local density | 1.895 | 1.780 | 1.370 | 1.000 |
| Aspect ratio | 0.604 | 0.735 | 0.761 | 0.824 |
| Separation | 0.729* | 0.769 | 0.622 | 0.742 |
| no_cofactor_tiering | | | | |
| Crossings/edge | 0.000 | 0.000 | 0.000 | 0.000 |
| Axis-aligned | 0.615 | 0.600 | 0.714 | 0.726 |
| Local density | 1.849 | 1.845 | 1.804 | 1.000 |
| Aspect ratio | 0.784 | 0.785 | 0.681 | 0.678 |
| Separation | 0.846 | 0.741 | 0.748 | 0.846 |
| no_fba | | | | |
| Crossings/edge | 0.000 | 0.000 | 0.000 | 0.000 |
| Axis-aligned | 0.515* | 0.583* | 0.714 | 0.667 |
| Local density | 1.807 | 1.813 | 1.833 | 1.000 |
| Aspect ratio | 0.817 | 0.869 | 0.800 | 0.876 |
| Separation | 0.846 | 0.824 | 0.783 | 0.846 |
| no_gap_filling | | | | |
| Crossings/edge | 0.000* | 0.000* | 0.000 | 0.000 |
| Axis-aligned | 0.610 | 0.600 | 0.716 | 0.714 |
| Local density | 1.867 | 1.882 | 1.802 | 1.875 |
| Aspect ratio | 0.653* | 0.568* | 0.559* | 0.571* |
| Separation | 0.679* | 0.456* | 0.492* | 0.688* |
| no_rings | | | | |
| Crossings/edge | 0.000 | 0.000 | 0.000 | 0.000 |
| Axis-aligned | 0.610 | 0.608 | 0.716 | 0.714 |
| Local density | 1.867 | 1.854 | 1.000 | 1.000 |
| Aspect ratio | 0.790 | 0.833 | 0.770 | 0.678 |
| Separation | 0.846 | 0.742 | 0.757 | 0.846 |

#### 6 Per-model layout results

Median per metric for each method and model: the per-model detail behind the layout-quality comparison in the manuscript.

#### 7 KEGG agreement across the corpus

The distribution plotted in the manuscript, as numbers. Pooled counts every scored reaction once, so the large well-annotated models dominate it; the median weights every model equally.

#### 8 KEGG agreement on the four most densely annotated models

The manuscript reports agreement pooled over the 104 scorable models. The four models with the largest curated intersection are given separately here, since they are the ones for which the per-reaction detail is densest.

### 9 Reproduction

```
pip install -e .[bench,test]
python scripts/fetch_bigg.py
python scripts/fetch_kegg.py --orgs eco hsa sce mmu

pytest                                   # regression suite
python -m src.bench.calibrate            # curated KEGG band
python -m src.bench.compare  --model iJO1366
python -m src.bench.ablate   --model iJO1366
python -m src.bench.correctness_all       # KEGG agreement, all models
python -m src.bench.correctness_control   # chance baseline
python -m src.bench.correctness_sensitivity  # stricter reference definitions
python -m src.bench.corpus                # metrics over every released map
python -m src.bench.scaling

python scripts/make_tables.py --check    # tables match the result files
python scripts/make_figures.py           # figures, at their placed width
```

| Method | Crossings/edge | Axis-aligned | Local density | Aspect ratio | Occupancy | Separation |
|---|---|---|---|---|---|---|
| RECON1, 129 maps | | | | | | |
| MetaCarto | 0.000 | 0.526 | 1.847 | 0.866 | 0.452 | 0.846 |
| DOT | 0.000 | 0.583 | 1.798 | 1.637 | 0.351 | 1.000 |
| NEATO | 0.000 | 0.000 | 1.789 | 1.038 | 0.415 | 0.738 |
| FDP | 0.035 | 0.000 | 1.706 | 1.003 | 0.421 | 0.773 |
| spring | 0.000 | 0.029 | 1.967 | 1.015 | 0.623 | 0.751 |
| KK | 0.174 | 0.000 | 1.913 | 0.989 | 0.526 | 0.355 |
| iAF692, 33 maps | | | | | | |
| MetaCarto | 0.000 | 0.786 | 1.822 | 0.697 | 0.389 | 0.769 |
| DOT | 0.000 | 0.786 | 1.826 | 0.923 | 0.305 | 0.846 |
| NEATO | 0.000 | 0.000 | 1.839 | 1.050 | 0.425 | 0.692 |
| FDP | 0.023 | 0.000 | 1.000 | 1.023 | 0.362 | 0.625 |
| spring | 0.000 | 0.000 | 1.792 | 1.059 | 0.623 | 0.763 |
| KK | 0.100 | 0.000 | 1.800 | 0.924 | 0.412 | 0.352 |
| iCHOv1, 214 maps | | | | | | |
| MetaCarto | 0.000 | 0.545 | 1.872 | 0.935 | 0.427 | 0.846 |
| DOT | 0.000 | 0.586 | 1.882 | 1.839 | 0.342 | 1.000 |
| NEATO | 0.000 | 0.000 | 1.822 | 1.005 | 0.427 | 0.727 |
| FDP | 0.000 | 0.000 | 1.793 | 1.020 | 0.402 | 0.739 |
| spring | 0.000 | 0.030 | 2.213 | 1.015 | 0.623 | 0.737 |
| KK | 0.168 | 0.017 | 1.916 | 1.001 | 0.522 | 0.367 |
| iJO1366, 93 maps | | | | | | |
| MetaCarto | 0.000 | 0.610 | 1.770 | 0.804 | 0.408 | 0.846 |
| DOT | 0.000 | 0.627 | 1.783 | 1.426 | 0.314 | 0.699 |
| NEATO | 0.000 | 0.000 | 1.784 | 1.006 | 0.410 | 0.698 |
| FDP | 0.020 | 0.000 | 1.000 | 1.014 | 0.406 | 0.679 |
| spring | 0.000 | 0.000 | 1.909 | 1.016 | 0.631 | 0.769 |
| KK | 0.231 | 0.018 | 1.852 | 0.956 | 0.524 | 0.367 |
| iMM904, 68 maps | | | | | | |
| MetaCarto | 0.000 | 0.600 | 1.803 | 0.806 | 0.404 | 0.731 |
| DOT | 0.000 | 0.613 | 1.031 | 1.528 | 0.324 | 0.846 |
| NEATO | 0.000 | 0.000 | 1.381 | 1.023 | 0.391 | 0.726 |
| FDP | 0.000 | 0.000 | 1.000 | 1.005 | 0.383 | 0.742 |
| spring | 0.000 | 0.021 | 1.894 | 1.021 | 0.623 | 0.760 |
| KK | 0.289 | 0.013 | 1.893 | 0.976 | 0.485 | 0.322 |
| iNJ661, 52 maps | | | | | | |
| MetaCarto | 0.000 | 0.714 | 1.811 | 0.770 | 0.401 | 0.757 |
| DOT | 0.000 | 0.725 | 1.250 | 1.016 | 0.309 | 0.552 |
| NEATO | 0.000 | 0.000 | 1.000 | 1.001 | 0.380 | 0.683 |
| FDP | 0.000 | 0.000 | 1.000 | 1.027 | 0.374 | 0.624 |
| spring | 0.000 | 0.000 | 1.900 | 0.992 | 0.617 | 0.752 |
| KK | 0.154 | 0.000 | 1.365 | 1.007 | 0.460 | 0.367 |
| iSB619, 38 maps | | | | | | |
| MetaCarto | 0.000 | 0.724 | 1.000 | 0.688 | 0.386 | 0.846 |
| DOT | 0.000 | 0.743 | 1.104 | 1.164 | 0.285 | 0.694 |
| NEATO | 0.000 | 0.000 | 1.000 | 0.999 | 0.412 | 0.759 |
| FDP | 0.000 | 0.000 | 1.000 | 1.006 | 0.382 | 0.738 |
| spring | 0.000 | 0.030 | 1.900 | 1.008 | 0.608 | 0.756 |
| KK | 0.118 | 0.000 | 1.000 | 1.091 | 0.444 | 0.314 |
| iYO844, 49 maps | | | | | | |
| MetaCarto | 0.000 | 0.726 | 1.000 | 0.678 | 0.381 | 0.846 |
| DOT | 0.000 | 0.737 | 1.000 | 1.549 | 0.277 | 0.880 |
| NEATO | 0.000 | 0.000 | 1.000 | 1.017 | 0.384 | 0.745 |
| FDP | 0.000 | 0.018 | 1.000 | 1.028 | 0.384 | 0.773 |
| spring | 0.000 | 0.009 | 1.923 | 1.016 | 0.629 | 0.773 |
| KK | 0.200 | 0.009 | 1.875 | 0.983 | 0.471 | 0.367 |

| Variant | Pooled | Median | Range |
|---|---|---|---|
| MetaCarto | 94.4 | 94.8 | 73.9–97.3 |
| − carrier tier | 89.3 | 89.5 | 70.5–91.3 |
| − cofactor handling | 58.9 | 59.8 | 38.5–69.6 |

| Model | Scored | Full | −tier | −cof. |
|---|---|---|---|---|
| iJO1366 | 673 | 95.1 | 90.5 | 59.1 |
| iMM904 | 559 | 96.1 | 91.2 | 52.8 |
| iYO844 | 450 | 94.7 | 88.4 | 58.9 |
| iAF692 | 284 | 92.6 | 88.7 | 52.5 |
